# Notch and Wnt signalling interact for proper prosensory and non-sensory domain formation

**DOI:** 10.1101/2025.11.24.690138

**Authors:** Neelanjana Ray, Sai Manoz Lingamallu, Arjun Guha, Raj K. Ladher

## Abstract

The prosensory domain (PSD) of the developing inner ear develops into the sensory hair cells, supporting cells, and neurons necessary for hearing and balance. Although Notch–Jag1 signalling is known to maintain Sox2 expression in this domain, the mechanisms by which Notch sustains prosensory identity have remained unclear. Here, we show that this function does not require canonical RBPJκ-dependent transcription. Using chick in ovo electroporation, Notch reporter assays, and conditional mouse mutants, we find that canonical Notch signalling is active during neurogenesis but not within the Sox2-positive PSD. Jag1 loss leads to a gradual loss of Sox2, yet RBPJκ mutants retain Sox2, indicating a non-canonical Notch pathway. We further reveal that the non-sensory determinant Lmx1a is regulated by Wnt/β-catenin signalling and that manipulating Wnt activity shifts the balance between Lmx1a and Sox2 expression. Finally, we show that NICD can directly interact with β-catenin in the cytoplasm, suggesting that NICD sequesters β-catenin to suppress Wnt activity and stabilise Sox2 in the PSD. Collectively, these findings support a model where prosensory maintenance occurs through non-canonical Notch signalling acting together with Wnt, forming a combined WNTCH regulatory module that coordinates sensory and non-sensory fate specification.

## INTRODUCTION

Hearing and balance are essential for how vertebrates interact with their environment. In birds and mammals, these functions are carried out by the inner ear, a complex epithelial labyrinth embedded within the temporal bone. The inner ear fulfils dual sensory roles via mechanosensory hair cells, which are arranged with supporting cells in the organ of Corti for auditory detection and within the maculae and cristae for vestibular function (Groves and Fekete, 2012; Wu and Kelley, 2012) (Groves & Fekete, 2012; Fritzsch et al., 2015). Signals from these hair cells are relayed through neurons of the cochleovestibular ganglion to auditory and vestibular regions of the brain (Appler and Goodrich, 2011)(Appler & Goodrich, 2011). In addition to these sensory epithelia, the inner ear includes non-sensory structures, such as the semicircular canals, endolymphatic duct and sac, that shape the biomechanical and ionic environment required for function (Fekete and Wu, 2002) (Fekete & Wu, 2002). Despite their structural and functional diversity, all of these components arise from the otic placode (Ladher et al., 2010; Nelson et al., 2025).

The otic placode is a thickened region of cranial ectoderm adjacent to the hindbrain that invaginates and closes to form the otic vesicle (Sai and Ladher, 2015; Tamilkumar et al., 2025). Within this vesicle, a ventral domain marked by Sox2 generates the neurosensory lineages of the inner ear, including sensory hair cells, supporting cells, and neurons of the cochleovestibular ganglion (Dabdoub et al., 2008; Gu et al., 2016). The first cells to differentiate from this region are the neurons, beginning around E9.5 in the mouse, when cells delaminate from the otic epithelium and migrate to form the ganglion (Raft et al., 2007). The neurogenic region arises from the anteromedial portion of the otocyst, which overlaps extensively with the anterior–ventral prosensory domain that expresses Sox2 and Jag1. Thus, both neurogenic progenitors and prosensory precursors originate within a shared Sox2- positive epithelial field (Adam et al., 1998; Cole et al., 2000; Kiernan et al., 2006a; Neves et al., 2011). Within this region, Notch–Delta signalling drives the selection of neuronal precursors: Delta-expressing cells delaminate and adopt neuronal identity, while neighbouring Notch-receiving cells remain as epithelial progenitors (Abello et al., 2007; Adam et al., 1998). Since Sox2 marks both populations before their divergence, maintaining it is essential for preserving neurosensory competence while enabling neuronal differentiation. This shared molecular origin means that neurogenesis and prosensory specification must be carefully balanced to ensure the correct allocation of cells to prosensory and proneuronal lineages.

While the molecular mechanisms that regulate Sox2 induction and maintenance in the prosensory domain are not fully understood, several signalling pathways have been implicated. In particular, Notch–Jag1 signalling sustains Sox2 expression through lateral induction (Brooker et al., 2006; Daudet et al., 2007). As the prosensory domain matures, Notch-dependent lateral inhibition subsequently guides the differentiation of hair cells and supporting cells (Daudet and Zak, 2020; Kiernan, 2013). Canonically, Notch signalling is mediated by ligand-induced cleavage of the receptor, releasing the Notch intracellular domain (NICD), which enters the nucleus and forms a transcriptional activation complex with RBPJκ (Recombination signal binding protein for immunoglobulin kappa J region) and MAML(Mastermind-like protein) to induce targets such as Hes/Hey genes (Bray, 2006). In the absence of NICD, RBPJκ instead recruits co-repressors such as Groucho/TLE to silence transcription (Oswald and Kovall, 2018). However, in the inner ear, loss of Jag1 or inhibition of NICD formation leads to a marked reduction of Sox2, whereas RBPJκ mutants retain Sox2 in the prosensory domain (Basch et al., 2011; Daudet et al., 2007; Kiernan et al., 2006a; Neves et al., 2011). This divergence indicates that prosensory maintenance requires a non-canonical, RBPJκ-independent mode of Notch signalling, the mechanism of which has remained unresolved.

Here, we investigate how non-canonical Notch signalling regulates Sox2. We show that NICD suppresses antagonistic Wnt signalling within the prosensory domain to stabilise Sox2 expression. We further demonstrate that the non-sensory determinant Lmx1a (Lmx1b in chick) is a Wnt target (Abello et al., 2007; Chizhikov et al., 2021; Koo et al., 2009), and that NICD directly interacts with β-catenin in the cytoplasm, indicating direct cross-talk between Notch and Wnt pathways. Together, these findings reveal a previously unrecognised mode of Notch signalling in the inner ear, in which NICD and β-catenin signalling intersect to balance prosensory maintenance, proneural development and non-sensory specification.

## MATERIAL AND METHODS

### Animals and mouse lines

All animal experiments were conducted in compliance with the Institutional Animal Ethics Committee guidelines. To capture the prosensory specification event, we used an early-expressing Cre driver line. The Six1-enh21-cre (From the lab of Dr. Shigeru Sato, RIKEN) mouse line carries a transgenic Cre recombinase under the control of enhancer 21 of the *Six1* gene. This enhancer is specifically active in the otic-epibranchial progenitor domain (OEPD) beginning at embryonic day 8.5 (E8.5), the precursor of the otic placode, and labels all subsequent lineages (Ono et al., 2014; Sato et al., 2018).

Conditional alleles (flox/flox) for *Rbpj*κ (RBRC01101), *Jag1* (JAX:010618), and β*- catenin* loss-of-function (JAX:004152) and β*-catenin* gain-of-function (MGI:1858008), were maintained as homozygotes. These animals were crossed with Six1-cre–positive mice to generate homozygous conditional mutants (or heterozygous mutants for β-catenin GoF). Embryos were genotyped for both the Cre transgene and the corresponding floxed allele.

### Electroporation in chicken embryos

Fertilised chicken eggs were obtained from the Central Poultry Development Organisation and Training Institute (Hessaraghatta, India) and incubated at 37°C in a humidified incubator for 52 hours to reach Hamburger and Hamilton stage 14 (HH14; Hamburger and Hamilton, 1951). Embryos were staged under a stereomicroscope, and a small window was made in the eggshell to access the embryo.

A DNA solution (plasmid DNA mixed with 30% sucrose and Fast Green dye) was microinjected into the otic cup using a fine glass capillary needle after carefully removing the extraembryonic membranes. A few drops of Ringer’s solution were then added over the embryo to improve conductivity. For electroporation, a positive electrode (cathode) was inserted beneath the embryo through a small hole made at the blunt end of the egg, and the negative electrode (anode) was gently placed on the solution above the otic vesicle. An electrical pulse of 12 V was applied to facilitate DNA uptake. Following electroporation, the window was sealed, and the eggs were incubated for 24 hours before embryo fixation.

For the Notch reporter assay, embryos were electroporated with a plasmid containing the Hes5 promoter driving destabilized nuclear eGFP, together with a constitutively active reporter plasmid (CAG promoter–driven mCherry), as previously described (Mann et al., 2017).

### Explant of chicken or mouse tissue for inhibitor experiments

The region of E10.5 mouse embryos or HH17 chicken embryos containing the otic vesicle was dissected in ice-cold Hank’s balanced salt solution (HBSS). Tissue pieces were embedded in a 1:1 mixture of Matrigel™ and DMEM in a 4-well dish and cured at 37 °C for 20 min. Culture medium (DMEM supplemented with N2 and penicillin) was then added together with either vehicle (DMSO) or the indicated inhibitors at the required concentration. Tissues were incubated for 20–24 h at 37 °C and subsequently collected for downstream processing.

### Immunofluorescence

#### Tissue Preparation and Cryosectioning

Collected tissues/embryos were fixed in 4% paraformaldehyde (PFA) for 1 h at room temperature (RT) and washed in 1× PBS. Samples were equilibrated in 15% sucrose (w/v) in 1× PBS overnight at 4 °C. Once the tissue sank, it was embedded in 7.5% gelatin prepared in 15% sucrose/1× PBS, oriented carefully, and solidified on dry ice. Blocks were stored at −20 °C. Cryosections (20–25 µm) were cut at −24 °C and collected on charged glass slides (Matsunami). Slides were stored at −20 °C and warmed to 37 °C immediately before immunofluorescence.

#### Immunofluorescence Staining

Slides were rinsed in warm 0.5% Triton in 1× PBS to dissolve/remove gelatin, then blocked in blocking buffer (10% heat-inactivated, filtered goat serum; 1% BSA; 0.5% Triton in 1× PBS) for 2 h at RT or overnight at 4 °C. Sections were incubated overnight at 4 °C with primary antibodies diluted 1:200 in blocking buffer (see Table 1). After thorough washes, secondary antibodies (see Table 1), DAPI, and phalloidin were diluted in blocking buffer and applied for 1 h at RT in the dark. Slides were washed and mounted with Fluoroshield mounting medium.

**TABLE 1:**
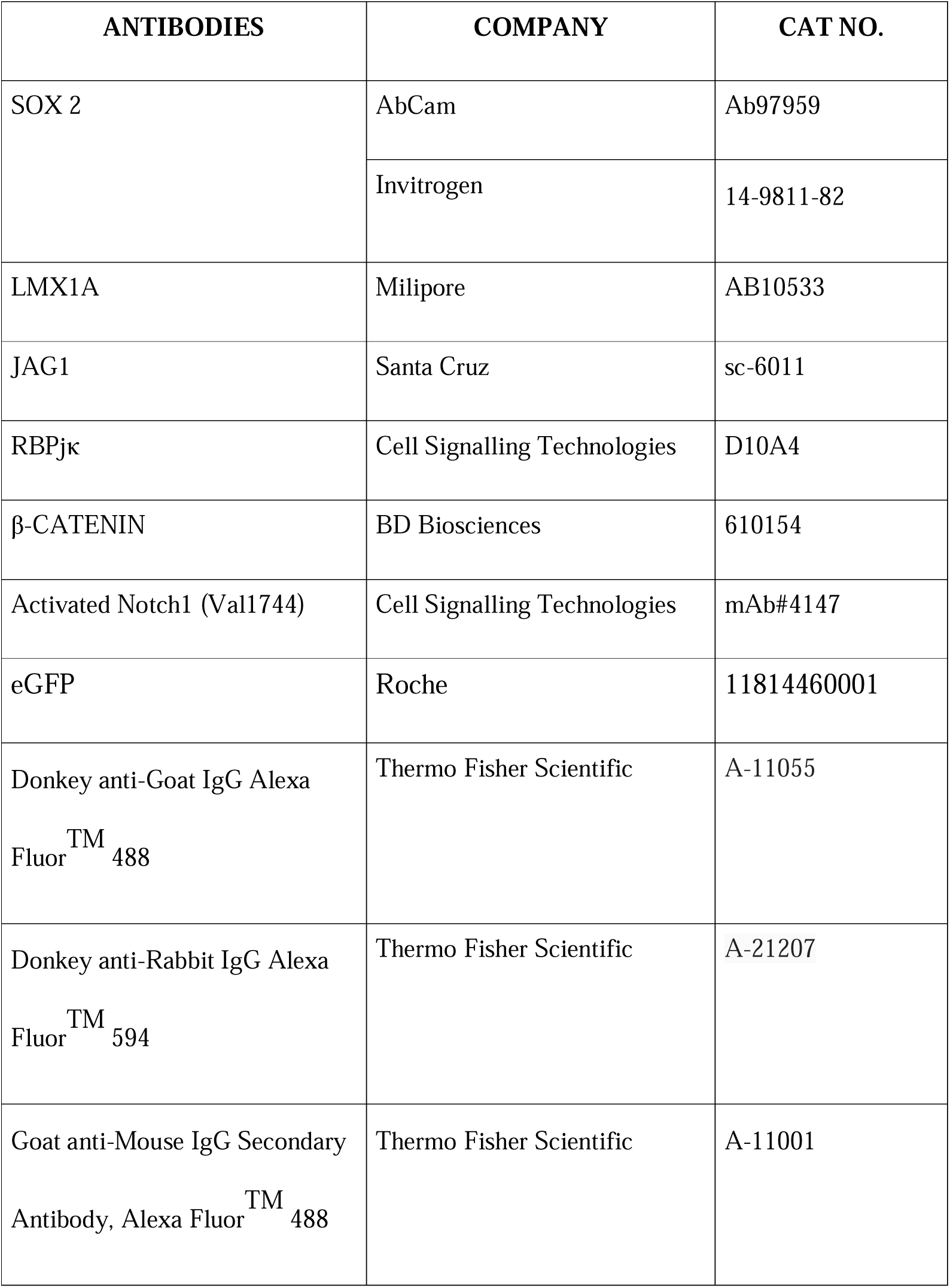
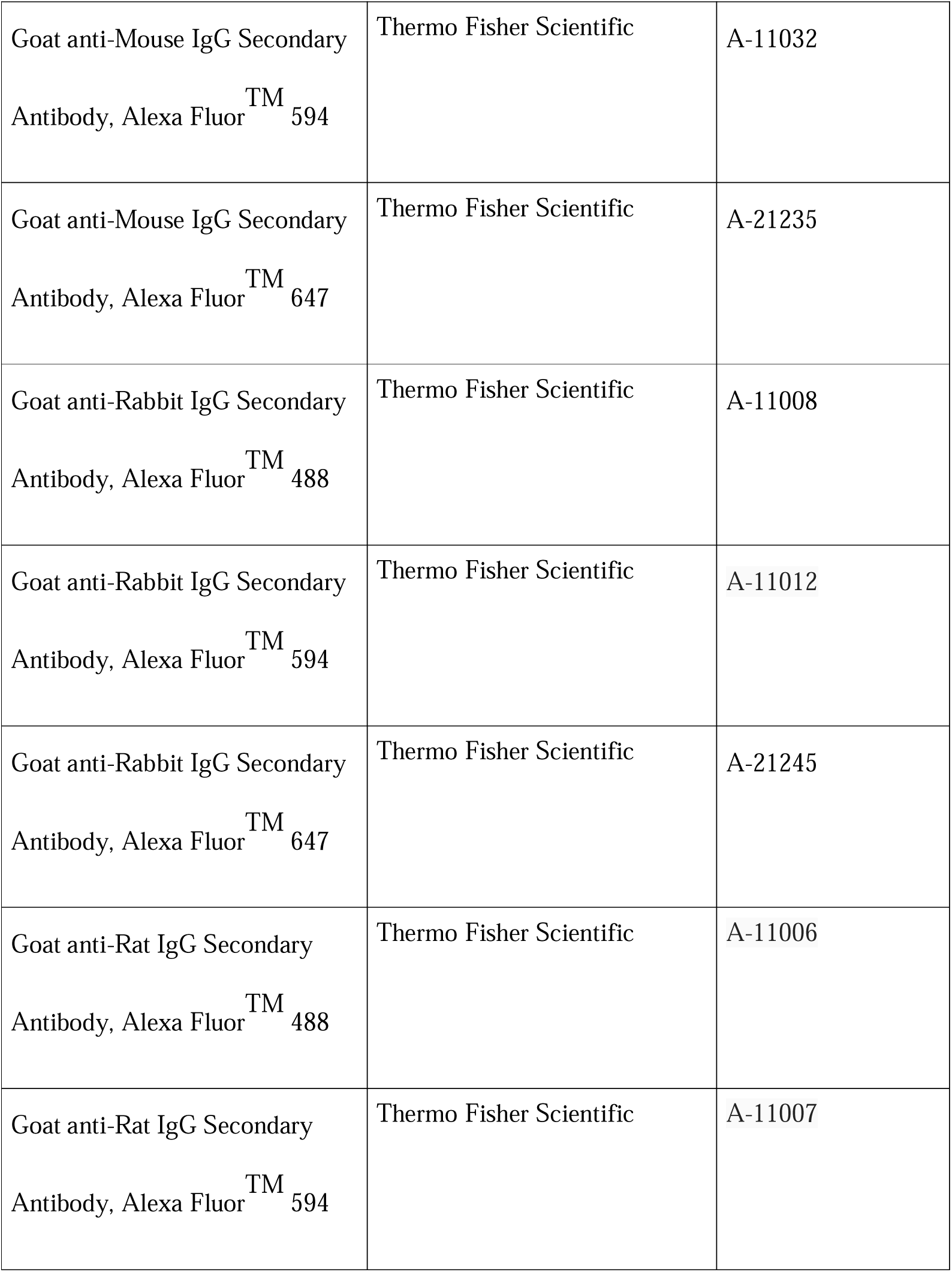
List of Antibodies.

#### Imaging and Processing

Confocal images were acquired on an Olympus FV3000 inverted microscope using FV31S- SW software. Raw image files were processed in ImageJ/Fiji.

#### Analysis of the cell population of various fates

Lmx1a expression was quantified in Fiji (ImageJ) by drawing a region of interest (ROI) across the otic vesicle on each section and generating intensity topographies with the 3D Surface Plot plugin. Based on the expression of Lmx1a and Sox2, four different cell populations (Sox2 positive, Lmx1a positive, High Sox2-Low Lmx1a, and High Sox2-High Lmx1a) were identified. The number and percentage of the cell population in the *Rbpj*κ mutant otic vesicle were determined using the cell counter plugin in Fiji (ImageJ).

### Co-immunoprecipitation

HEK293T cells were maintained under standard culture conditions (in Dulbecco’s Modified Eagle Medium (DMEM) supplemented with 10% Fetal Bovine Serum (FBS) at 37°C in 5% CO_2_) and transfected with the plasmids (see Table 2) using Mirus LTi reagent according to the manufacturer’s protocol. Cells were harvested 48 h post-transfection, lysed in lysis buffer, and processed for co-immunoprecipitation (co-IP).

**TABLE 2:**
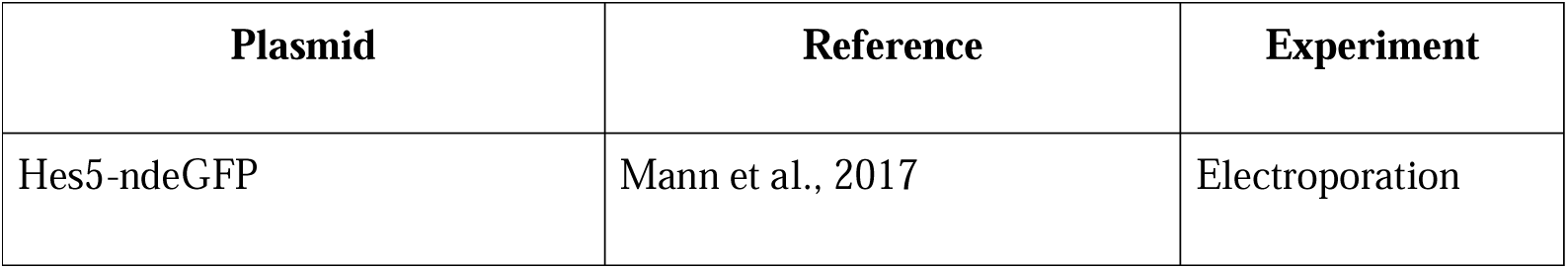
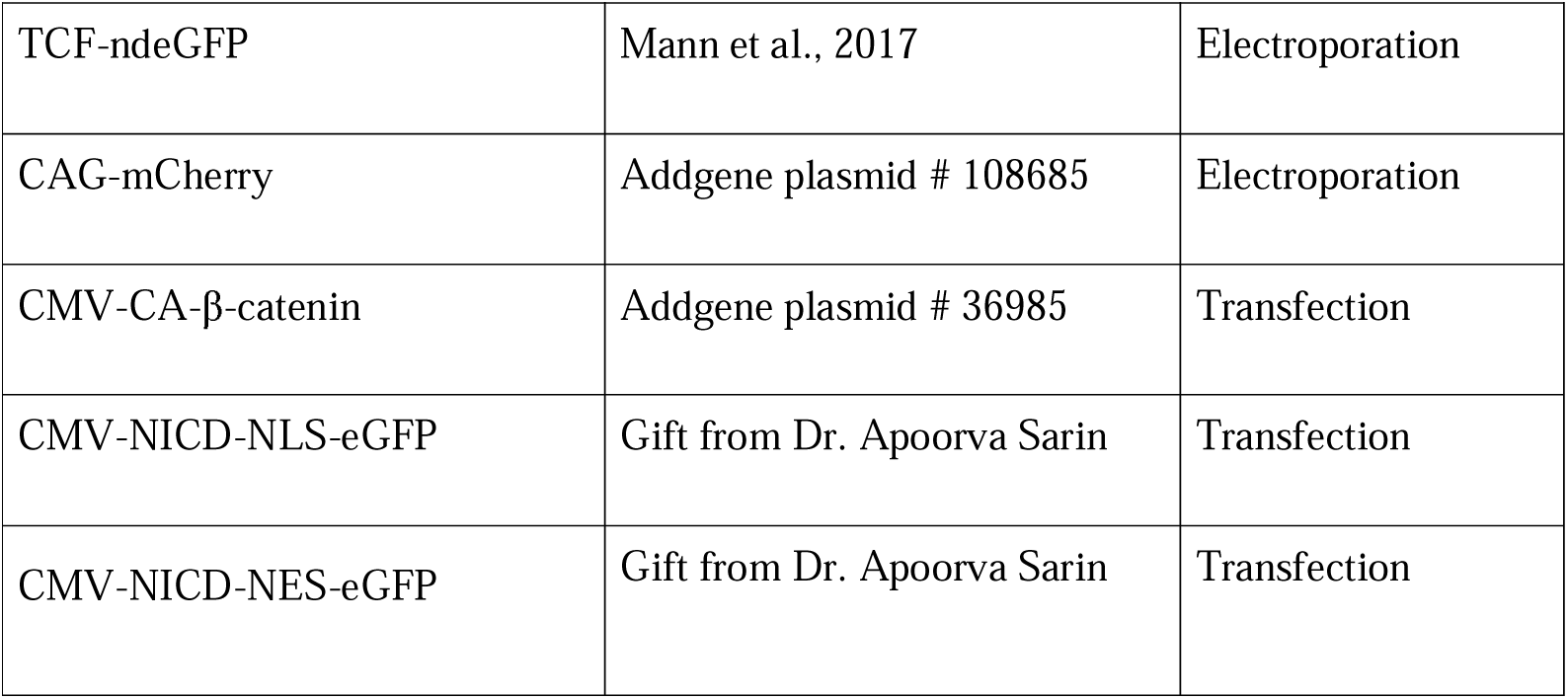
List of Plasmids.

| Plasmid | Reference | Experiment |
| --- | --- | --- |
| Hes5-ndeGFP | Mann et al., 2017 | Electroporation |
| TCF-ndeGFP | Mann et al., 2017 | Electroporation |
| CAG-mCherry | Addgene plasmid # 108685 | Electroporation |
| CMV-CA-β-catenin | Addgene plasmid # 36985 | Transfection |
| CMV-NICD-NLS-eGFP | Gift from Dr. Apoorva Sarin | Transfection |
| CMV-NICD-NES-eGFP | Gift from Dr. Apoorva Sarin | Transfection |

Cell or tissue lysates were incubated with magnetic Dynabeads conjugated to anti–β-catenin antibody under gentle rotation at 4 °C overnight. The supernatant was collected and analysed by SDS–PAGE and immunoblotting. Protein samples were mixed with loading buffer, resolved on 10% SDS–PAGE gels, and transferred onto PVDF membranes.

Membranes were probed with antibodies against active β-catenin (1:1500) and eGFP (1:5000), noting that the NICD–NES fusion protein is eGFP-tagged. HRP-conjugated secondary antibodies were applied, and chemiluminescent signals were detected using an ECL detection system.

### Paint filling

The paint-filling protocol was kindly provided by Dr. Doris Wu. Briefly, heads from E15.5 mouse embryos were fixed overnight in Bodian’s fixative, followed by dehydration in 100% ethanol for 2 days to ensure complete dehydration. Samples were then incubated in methyl salicylate until fully cleared.

Cleared heads were bisected along the midline, and the brain was carefully removed to expose the inner ear. White enamel paint, diluted in methyl salicylate, was injected into the mid-turn of the organ of Corti using a sharp glass capillary. In cases where the cochlea was not identifiable, injections were performed through the vestibular apparatus.

## RESULTS

### Notch signalling is required for Sox2 maintenance in PSD

The sensory cells of the inner ear arise from the prosensory domain (PSD), which is marked by Sox2 expression. At E8.5 in the mouse, the PSD is located on the ventral side of the otic vesicle and is established through the integration of multiple signalling pathways, including FGF, Notch, BMP, Wnt, and Shh (Dabdoub et al., 2008; Jacques et al., 2012; Kiernan et al., 2005; Ohyama et al., 2010; Riccomagno et al., 2002; Wright and Mansour, 2003; Zak and Daudet, 2021). These pathways act in a finely balanced manner, with some instructing and others repressing sensory fate, through extensive cross-regulation. Among them, Notch-mediated lateral induction, through its engagement with Jag1, has been shown to play an essential role in regulating Sox2 expression in the PSD (Brooker et al., 2006; Daudet et al., 2007; Kiernan et al., 2006b). Jag1 is expressed in the same region as the PSD (Supp Fig. 1A). Similar to already published data (Daudet et al., 2007), we also find that blocking the formation of NICD by inhibiting the activity of γ-secretase reduces the Sox2 expression in PSD (Fig. 1A–B).

**FIGURE 1:**
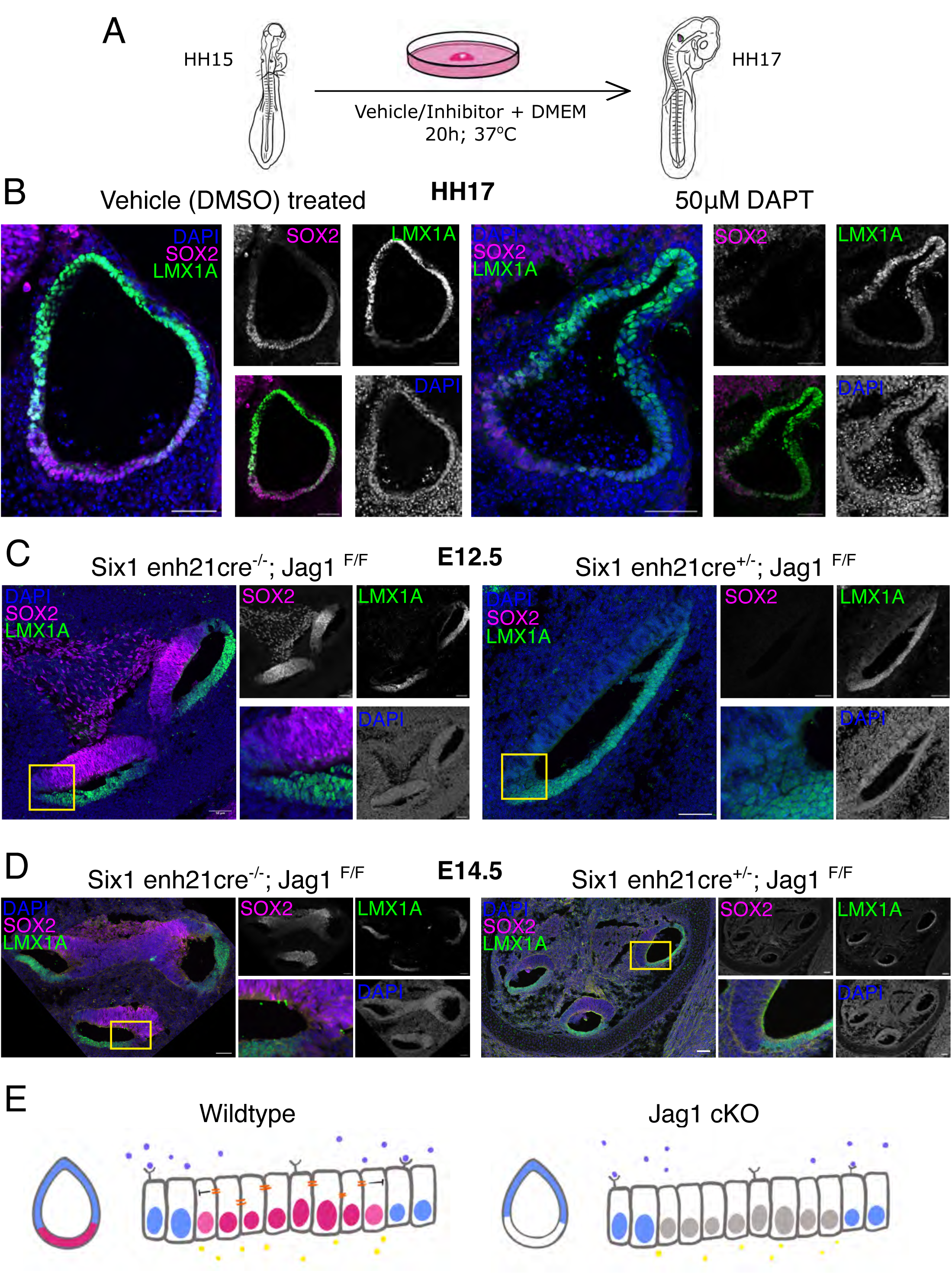
Sox2 expression depends on Jag1-mediated Notch signalling in the prosensory domain. (A) Experimental outline of explant experiments for checking the effect of the small molecule GSK3β inhibitor, DAPT, on HH15 chicken otocyst. (B) Dorso-ventral cross-section of the otic vesicle shows the expression of sensory determinant, Sox2 (in magenta), decreased in the DAPT-treated sample as compared to the DMSO-treated otocyst. The pattern of non-sensory determinant, Lmx1a remains similar. (C) E12.5 *Jag1* cKO embryo shows a decrease in Sox2 expression in the PSD, while Lmx1a is expressed in the non-sensory domain, like the wild type littermate (D) Sox2 expression remains very low to almost none in the sensory epithelium of E14.5 *Jag1* cKO inner ear, while the wildtype inner ear has well segregated sensory and non- sensory regions. All images have a scale bar of 50 μm. (E) Schematic summarising the phenotype of *Jag1* cKO epithelium. Yellow circles: endoderm-derived FGF-signalling proteins, violet circles: dorsalizing Wnt signal, orange lines: juxtacrine lateral induction mediated by Notch signalling, pink nuclei: Sox2 positive sensory lineage, purple nuclei: Lmx1a positive non-sensory lineage, and grey: Neither Sox2 nor Lmx1a positive cell population.

The expression of Sox2 in the otic vesicle is complementary to that of the LIM homeodomain transcription factor Lmx1a. In the chick, Lmx1a expression represses Sox2 during the partitioning of the PSD to different sensory structures within the otocyst (Mann et al., 2017). We thus asked if the down-regulation of Sox2 resulting from early Notch signalling perturbations could be due to the mis-regulation of Lmx1a expression. To do this, we deleted *Jag1* using the Six1-21-Cre driver, which is active in the otic placode from E8.5 in mice. In these mutants, Sox2 expression was detected in the otic epithelium until E11.5 but was down-regulated and undetectable by E12.5. Interestingly, by E14.5, faint Sox2 expression reappeared (Fig. 1C–D), possibly reflecting compensatory activation by other signalling pathways such as FGF (Hayashi et al., 2007). To determine whether the loss of Sox2 resulted from a change in PSD fate, we examined Lmx1a expression in these mutants. Lmx1a remained restricted to the non-sensory domain, similar to wild-type controls. These findings indicate that Jag1-dependent Notch signalling is required for the maintenance of Sox2 in the PSD but does not influence Lmx1a expression or non-sensory lineage specification.

### SOX2 and LMX1A —dual positive cell population emerge in the absence of RBPjκ

Previous studies have conflicting conclusions on whether canonical Notch signalling is required for Sox2 expression in the PSD (Basch et al., 2011; Yamamoto et al., 2011). To determine whether canonical Notch signalling is active in the putative PSD, we first examined the chick model. In the chick, expression of Sox2 in the otic vesicle is complementary to that of Lmx1a, a pattern that is conserved in the mouse. Furthermore, Sox2 regulation in chick is also controlled by Notch-dependent lateral induction (Neves et al., 2011). To examine whether Sox2 in the PSD is regulated by canonical Notch signalling, we asked if inhibition of Rbjκ activity altered Sox2 expression. Using an explant approach, we treated the otic vesicle with a small molecule inhibitor of RBPjκ- RIN1. We observed that Sox2 expression remained unchanged, indicating an RBPjκ-independent mode of Sox2 expression in the otic vesicle (Fig. 2A–B).

**FIGURE 2:**
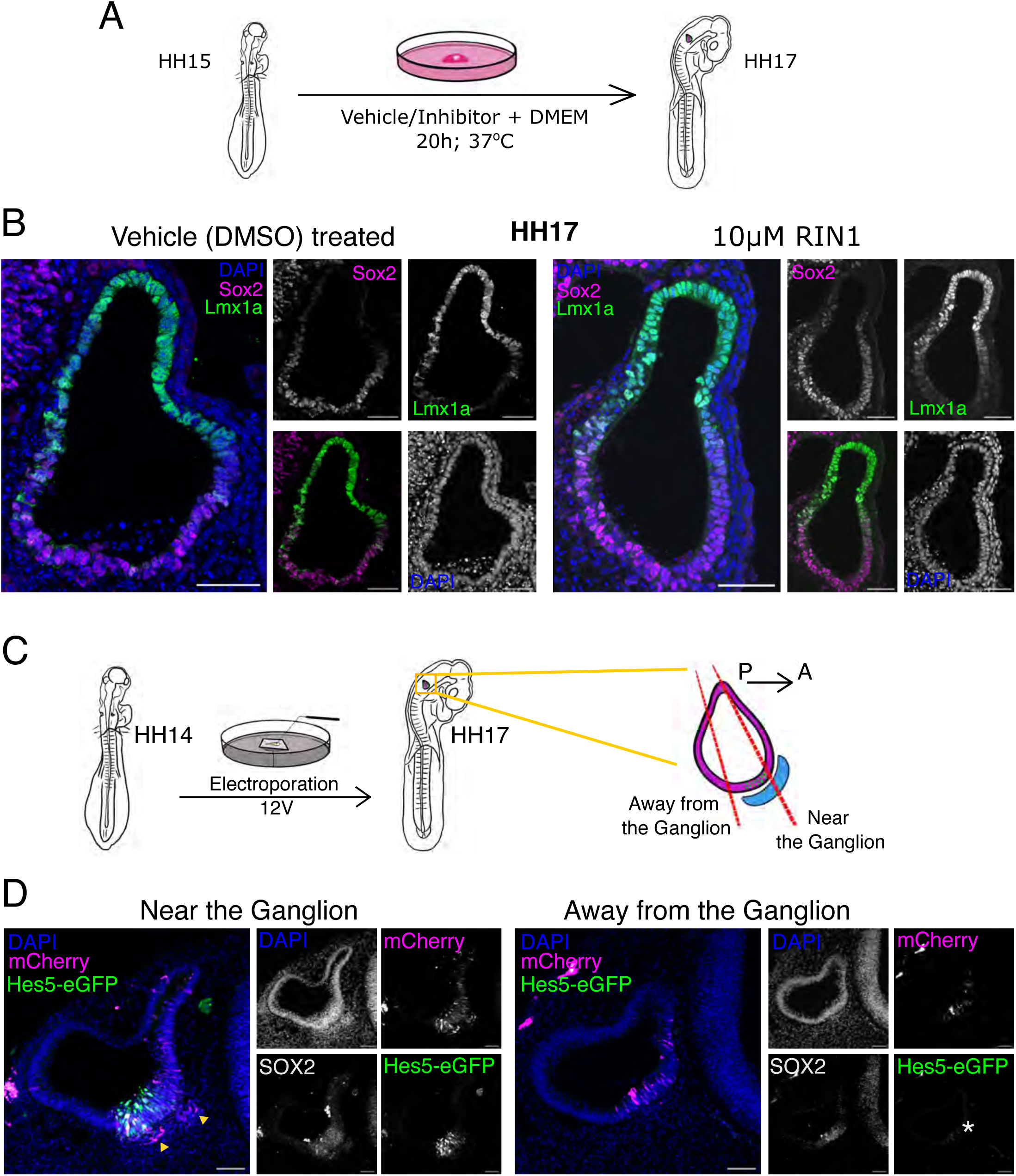
Non-canonical Notch signalling maintains Sox2 expression in PSD. (A) Experimental outline of *in vitro* explant for checking the effect of small molecule RBPjκ inhibitor, RIN1, on HH15 chicken otocyst. (B) Inhibitor of RBPjκ- RIN1 does not change Sox2 or Lmx1a expression in the explant culture of HH17 otocyst. (C) Schematic showing the experimental design of assaying for canonical notch reporter activity in the chicken otic vesicle. Electroporation was done at 12 Volts. 20um-thick sections were collected based on proximity to the ganglion. (D) PSD near the ganglion of the HH17 chick embryo (yellow arrowhead marking the delaminated neurons) shows activity of the Hes5-eGFP reporter (green) in the regions electroporated (marked by magenta-mCherry) and expressing Sox2 (grey), whereas low to no activity of the Notch reporter (marked by white asterisk) is observed away from the ganglion.

We next employed a Hes5-GFP reporter construct as an indicator of pathway activity (Mann et al., 2017). This encodes as destabilised nuclear eGFP driven by Hes5 regulatory sequences. The plasmid was electroporated into the otic vesicle of HH14 chicken embryos. By HH17, reporter expression was observed in delaminating neurons, consistent with previous studies indicating that neuronal specification depends on canonical Notch signalling (Abello et al., 2007). Nonetheless, eGFP was not detected in Sox2-positive epithelial cells of the PSD, despite successful electroporation confirmed by the expression of the mCherry tracer (Fig. 2C–D). These results suggest that canonical Notch signalling is not active in the PSD at this stage.

To understand how the dynamics of Sox2 expression are influenced by canonical Notch signalling, we next examined both Sox2 and Lmx1a expression in mouse otocysts from conditional *Rbpj*κ mutants. Immunostaining with an anti-RBPjκ antibody confirmed the loss of protein in mutant vesicles from E10.5 (Supp Fig. 1B–C). Despite this loss, Sox2 expression at E10.5 was similar to that of wild-type and heterozygous controls, both in intensity and spatial distribution (Fig. 3A). Interestingly, Lmx1a expression was altered. In control embryos, Sox2 and Lmx1a occupy largely reciprocal domains, with only a small boundary region of overlap where Lmx1a levels are low and Sox2 expression is high. In *Rbpj*κ mutants, the Lmx1a domain expanded considerably, creating a broad overlap with Sox2 and resulting in an increased number of Lmx1a-high/Sox2-high dual-positive cells (Fig. 3B–C). By E14.5, this overlap resolved in the cochlear duct, with marker expression returning to a near-normal pattern, although Lmx1a expression remained abnormal in the vestibular organs (Fig. 3D).

**FIGURE 3:**
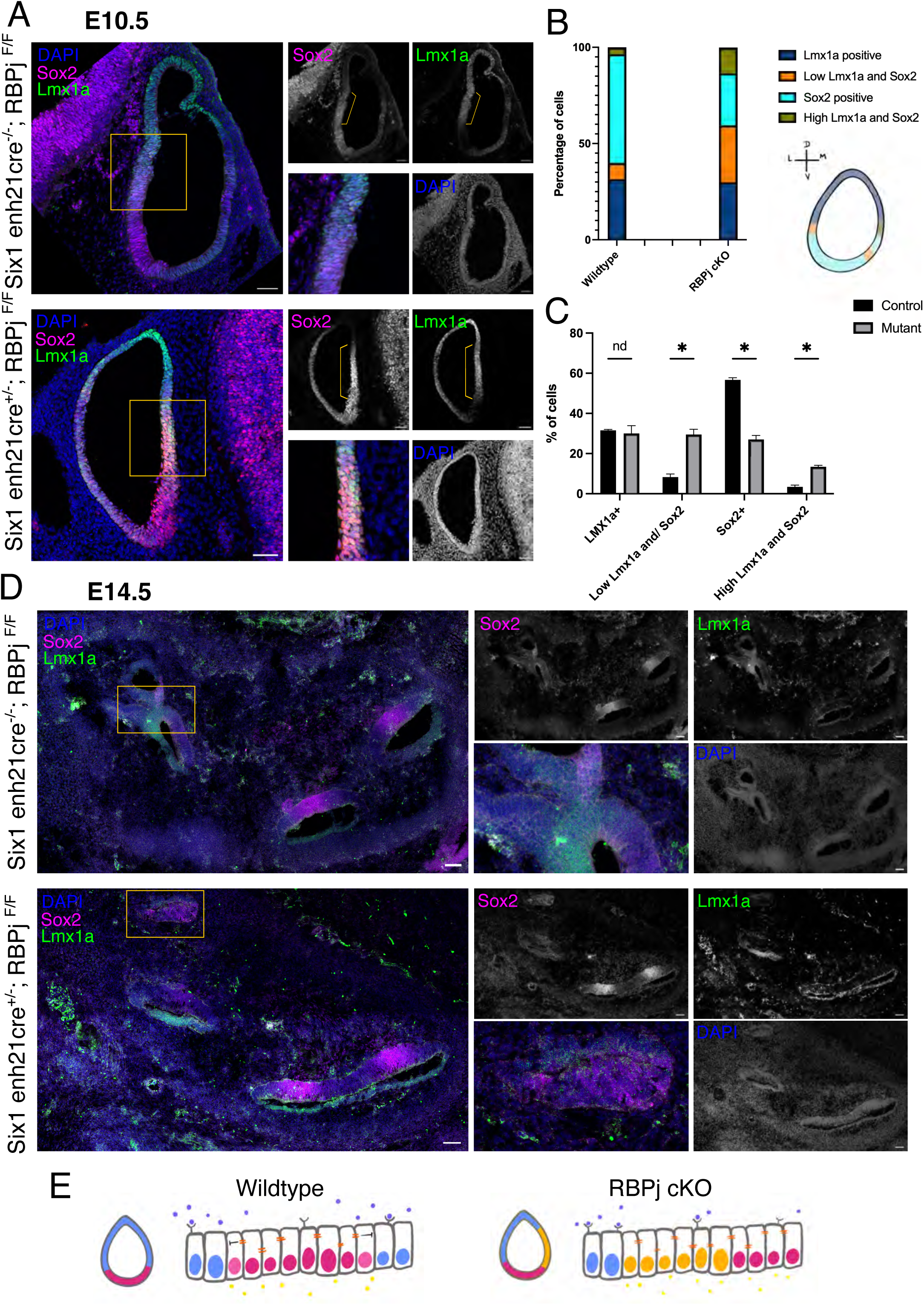
*Rbpj*κ mutant shows overlapping SOX2 and LMX1A expressing region. (A) Otic vesicle at E10.5 shows distinct Sox2 (magenta) and Lmx1a (green) expressing region, which starts overlapping in the *Rbpj*κ conditional knockout embryo. (B) The percentage of cells in four different cell-fate lineages is plotted for *Rpbj*κ cKO versus wild type. (C) Change in the number of cells expressing only Sox2, and the population of dual-positive cells is significant, with respective p value 0.000283 (low Lmx1a and Sox2), 0.000101 (high Lmx1a and Sox2) and 0.000025 (Sox2 expressing). (D) In E14.5, the *Rbpj*κ cKO has reduced Lmx1a-positive cells in the developing vestibular apparatus. All images have a scale bar of 50 μm. (E) Schematic to summarise the phenotype of *Rbpj*κ cKO epithelium. Yellow circles: endoderm-derived FGF-signalling proteins, violet circles: dorsalizing Wnt signal, orange lines: juxtacrine lateral induction mediated by Notch signalling, pink nuclei: Sox2 positive sensory lineage, purple nuclei: Lmx1a positive non-sensory lineage, and mustard yellow: dual positive cell lineage.

These results suggest that although the non-sensory marker Lmx1a is not directly controlled by Jag1-mediated Notch signalling, its expression is affected by the loss of RBPjκ. Furthermore, Sox2 expression in the PSD is largely independent of canonical Notch signalling, however cross-regulation between signalling pathways could influence domain boundaries and cellular identity during otic development.

### Expression of LMX1A is controlled by Wnt signalling

To investigate whether signalling pathway interactions regulate Sox2 expression in the PSD, we next examined how Lmx1a expression is controlled. Wnt signalling has been implicated in regulating Lmx1a during the development of midbrain dopaminergic neurons (Chung et al., 2009). Moreover, it is known to play a role in dorsal patterning of the inner ear (Noda et al., 2012; Riccomagno et al., 2005). We therefore explored the regulation of Lmx1a by Wnt signalling.

We employed the chick system to investigate Wnt activity using the inhibitor XAV939 and the agonist CHIR99021. XAV939, a tankyrase inhibitor, stabilises Axin, leading to reduced β-catenin accumulation (Huang et al., 2009). Treatment of HH15 otocysts with XAV939 for 20 hours resulted in decreased β-catenin levels. In these explants, we observed a downregulation of Lmx1a expression (Fig. 4A). Conversely, CHIR99021 inhibits GSK3β, which normally promotes β-catenin degradation (An et al., 2010). Treatment with CHIR99021 led to β-catenin upregulation and widespread Lmx1a expression not only within the otocyst but also in surrounding tissues (Fig. 4B). These findings indicate that Lmx1a is regulated by Wnt signalling.

**FIGURE 4:**
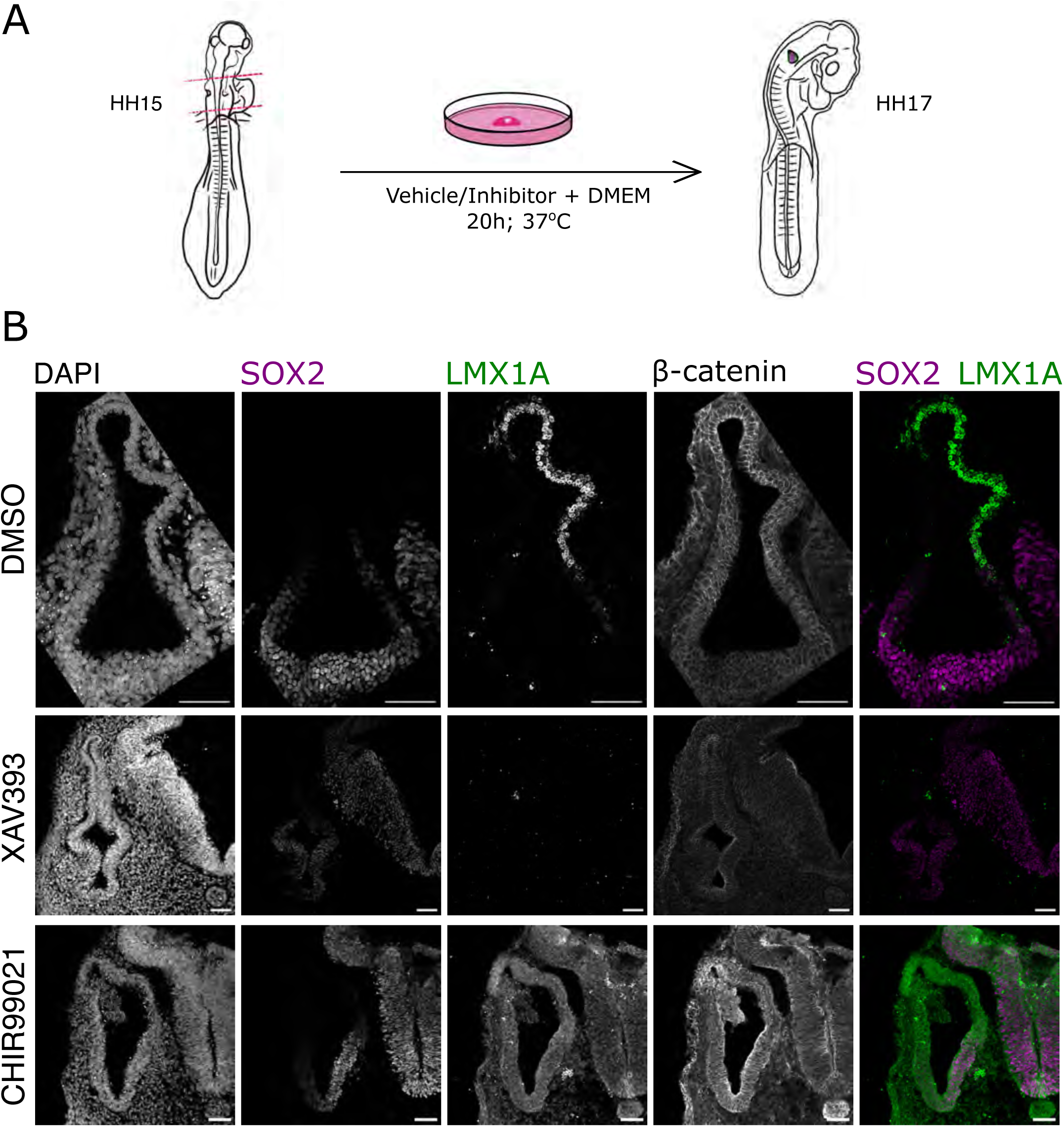
Lmx1a expression is modulated by a small molecule inhibitor of GSK3β *ex vivo*. (A) Experimental design of the explant culture using HH15 chicken embryo. (B) Expression of both Sox2 and Lmx1a is downregulated in XAV-treated Wnt inhibition conditions from their respective compartments. Overactivating Wnt using GSK3β inhibition causes diffused expression of Lmx1a and a very slight decrease in Sox2 expression. All images have a scale bar of 50 μm.

To further confirm this regulatory relationship, we used genetic mouse models to manipulate Wnt signalling. In the absence of Wnt activity, β-catenin is targeted for proteasomal degradation by a sequence encoded by exon 3 of the Ctnnb1 gene. We first used a β-catenin gain-of-function (GOF) mouse model, in which exon 3 is conditionally removed, resulting in stabilised β-catenin (Harada et al., 1999). This line was crossed with Six1-enh21- Cre to activate ectopic Wnt signalling in the inner ear. During early otocyst patterning (E9.5), Sox2 showed discontinuous patches of expression on the ventral side of the otocyst, while Lmx1a was expressed broadly throughout the otocyst and in the neural tube (Fig. 5A). By E13.5, sections revealed Lmx1a and Sox2 expression in structures likely corresponding to vestibular sensory organs (Fig. 5B). However, the cochlea appeared to be absent, a finding confirmed by paint-fill preparations at E15.5 (Fig. 5C).

**FIGURE 5:**
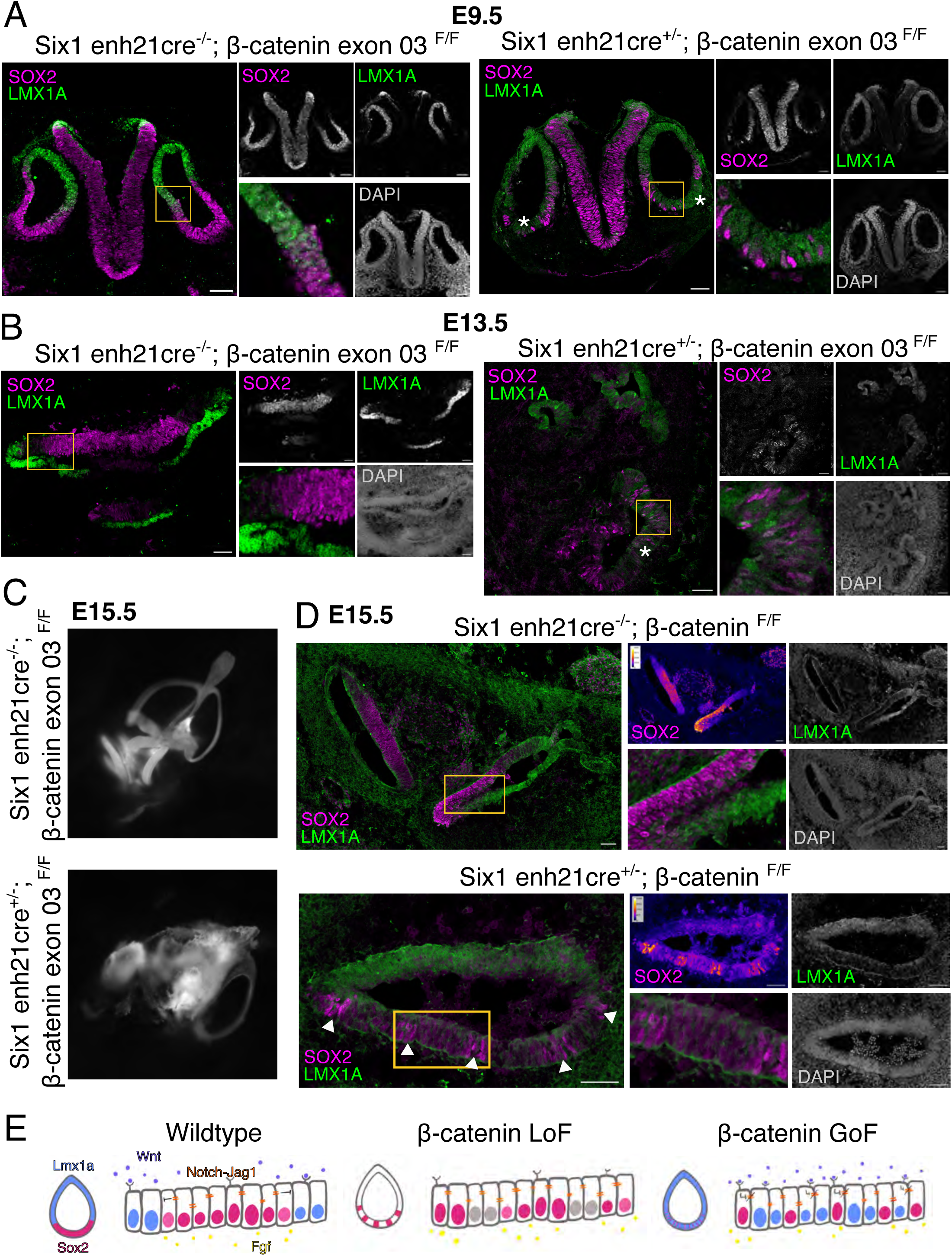
Moderate levels of β-catenin mediated Wnt signalling are required for the proper size and pattern of PSD. (A) Transverse section view of the thoracic region of the E9.5 mouse embryo showing the sensory and non-sensory domains marked by Sox2 (magenta) and Lmx1a (green) in wildtype and β-catenin gain-of-function mutant. Mutant shows an absence of proper PSD size marked by the white asterisk. (B) By E13.5 β-catenin gain-of-function mutant has defective morphogenesis of the cochlear tube, which has unclear morphology as well as improperly segregated sensory and non-sensory domains. (C) Morphology of the E15.5 β-catenin gain-of-function mutant versus the wildtype visualised by paintfilling the inner ear canal. (D) Pattern of the sensory- non sensory domains in E15.5 wildtype and β-catenin loss-of-function mutant. All images have a scale bar of 50 μm. (E) Schematic to summarise the β-catenin gain-of-function and loss-of-function mutant phenotypes.

To investigate the requirement for Wnt signalling in regulating Lmx1a and Sox2, we examined β-catenin loss-of-function (LOF) mutants where β-catenin was removed from the inner ear. In E9.5 otocysts from these mutants, Lmx1a expression was significantly downregulated, and Sox2 expression shifted towards the dorsomedial region. By E13.5, the mutant cochlea showed patches of Sox2 expression, with low levels of Lmx1a detected in the roof of the cochlear duct (Fig. 5D).

Taken together, these findings show that Lmx1a expression is regulated by Wnt signalling. Furthermore, perturbations in Wnt signalling alter Sox2 expression, indicating cross-regulation between the signalling pathways that establish prosensory and non-sensory domains of the inner ear.

### NICD and β-catenin can interact *in vitro* and bind to each other

Given the interdependence of Sox2 and Lmx1a on Notch and Wnt signalling, we hypothesised that there may be direct cross-talk between the two pathways. To test this, we examined the interaction between the key intracellular mediators of the pathways, NICD and β-catenin. We transfected HEK293T cells with constitutively active β-catenin (CA–β- catenin) and one of two NICD constructs: one with a nuclear localisation sequence (NICD- NLS-eGFP) and one with a nuclear export sequence, which is located in the cytoplasm (NICD-NES-eGFP) (Fig. 6A). Co-immunoprecipitation using a β-catenin antibody, followed by Western blotting for eGFP, revealed that β-catenin and NICD-NES-eGFP interact (Fig. 6B). In contrast, NICD-NLS-eGFP does not interact with β-catenin. These data suggest that in cells where both signals are active, NICD and β-catenin may be sequestered in the cytoplasm, limiting β-catenin mediated Wnt signalling and thus stabilising Sox2 expression in the PSD.

**FIGURE 6:**
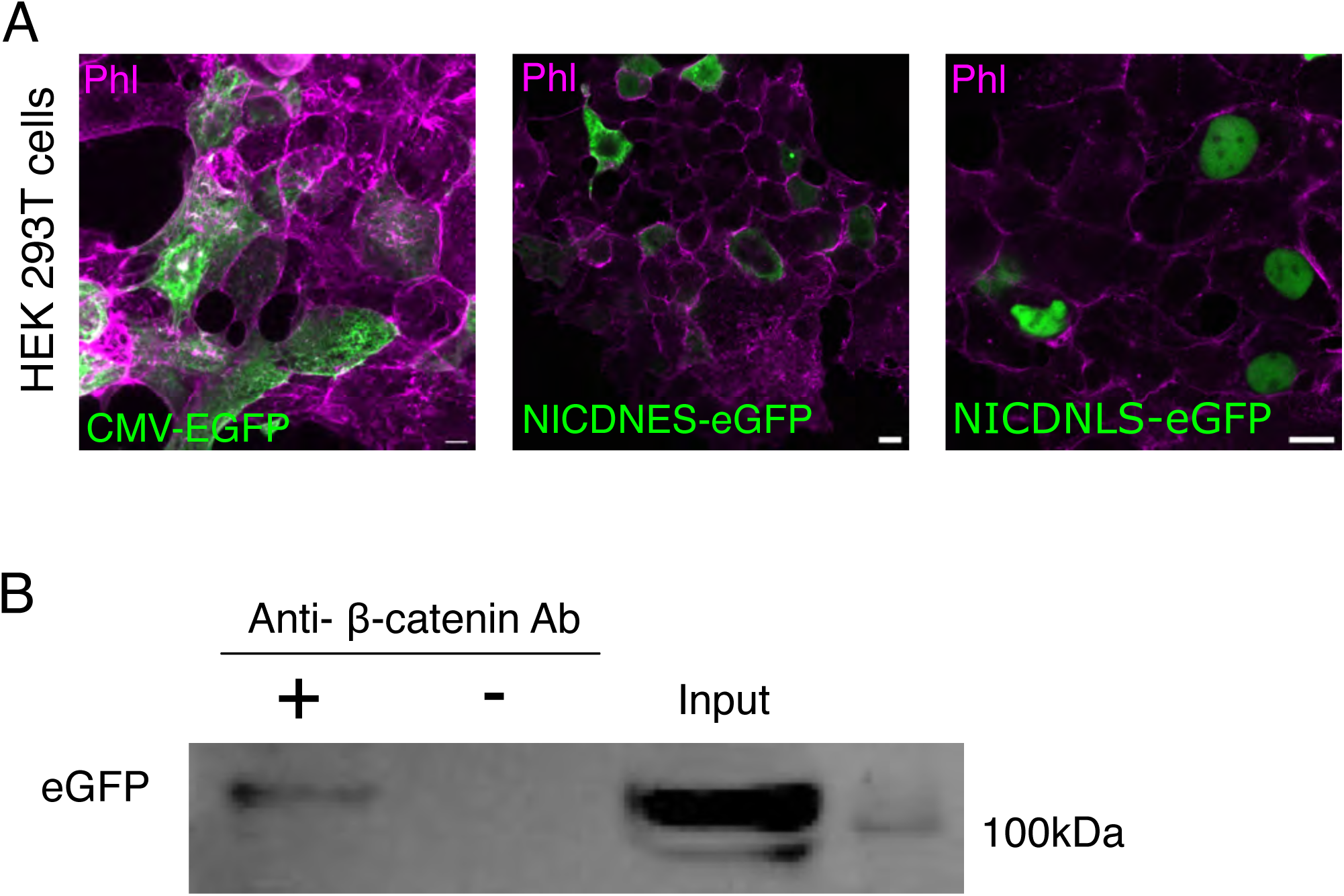
β-catenin and NICD biochemically interact with each other in the cytoplasm. (A) Immunofluorescence showing the transfected plasmid variant- CMV-EGFP throughout the cell, NICDNES-EGFP in the cytoplasm and NICDNLS-EGFP in the nucleus by staining for EGFP (green) along with F-actin stained by phalloidin (magenta). All images have a scale bar of 5μm. (B) Western blot analysis to show the presence of NICDNES-EGFP bound to the β-catenin soaked DynaBead. The calculated molecular size of β-catenin is 86kDa; however observed band is slightly more than 100kDa, probably due to post-translational modification.

### Removing JAG1 in an over-activated Wnt signalling condition disrupts the dorso-ventral axis of the otic vesicle

Our data indicate that Wnt signalling and Notch/Jag1 signalling interact to regulate prosensory patterning. To examine how this cross-regulation shapes the expression domains of Lmx1a and Sox2, we generated double mutants in which both pathways were perturbed by combining Jag1 conditional knockout (CKO) with β-catenin gain-of-function (GOF). At E9.5, Lmx1a expression was detected throughout the otic epithelium and Sox2 expression was observed in discontinuous patches (Fig. 7A). This was unlike in the Jag1 mutant but was similar to the β-catenin GOF mutant; however, these patches appeared in both dorsal and ventral regions, with Sox2 expression levels in the dorsal patches notably lower than in the ventral regions. In a β-catenin GOF: Jag1 heterozygous mutant, the expression of discontinuous Sox2 was restricted only to the ventral half. By E11.5, Sox2 expression was no longer detectable, and Lmx1a remained broadly expressed throughout the inner ear (Fig. 7B).

**FIGURE 7:**
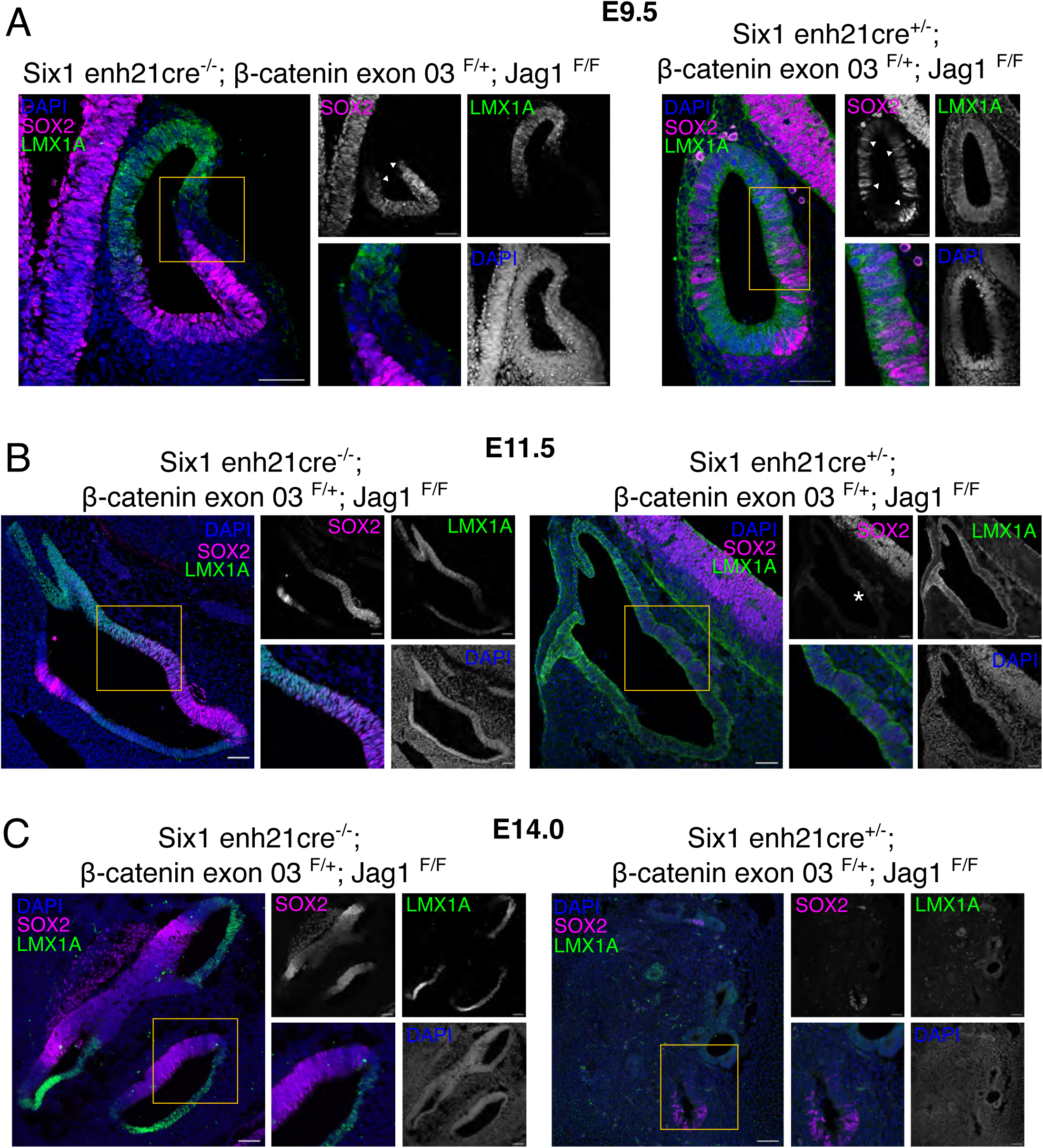
Disrupting Jag1 and overactivating the Wnt signalling leads to complete abrogation of Dorso-Ventral polarity. (A) In the E9.5 embryo, Sox2 (magenta) expression becomes disrupted, and such cells are present throughout the otic vesicle, interspersed with Lmx1a-positive cells (green) in the double mutant embryo, β-catenin gain-of-function, and *Jag1* cKO, instead of properly segregated sensory and non-sensory domains. (B) Transverse section of E11.5 inner ear shows complete loss of Sox2 expression in the double mutants as compared to the wildtype. (C) In E14.0 inner ear, immunofluorescence shows the sparse presence of both the sensory (Sox2 positive) and the non-sensory (Lmx1a positive) cell lineage in the double mutant. All images have a scale bar of 50μm.

**FIGURE 8:**
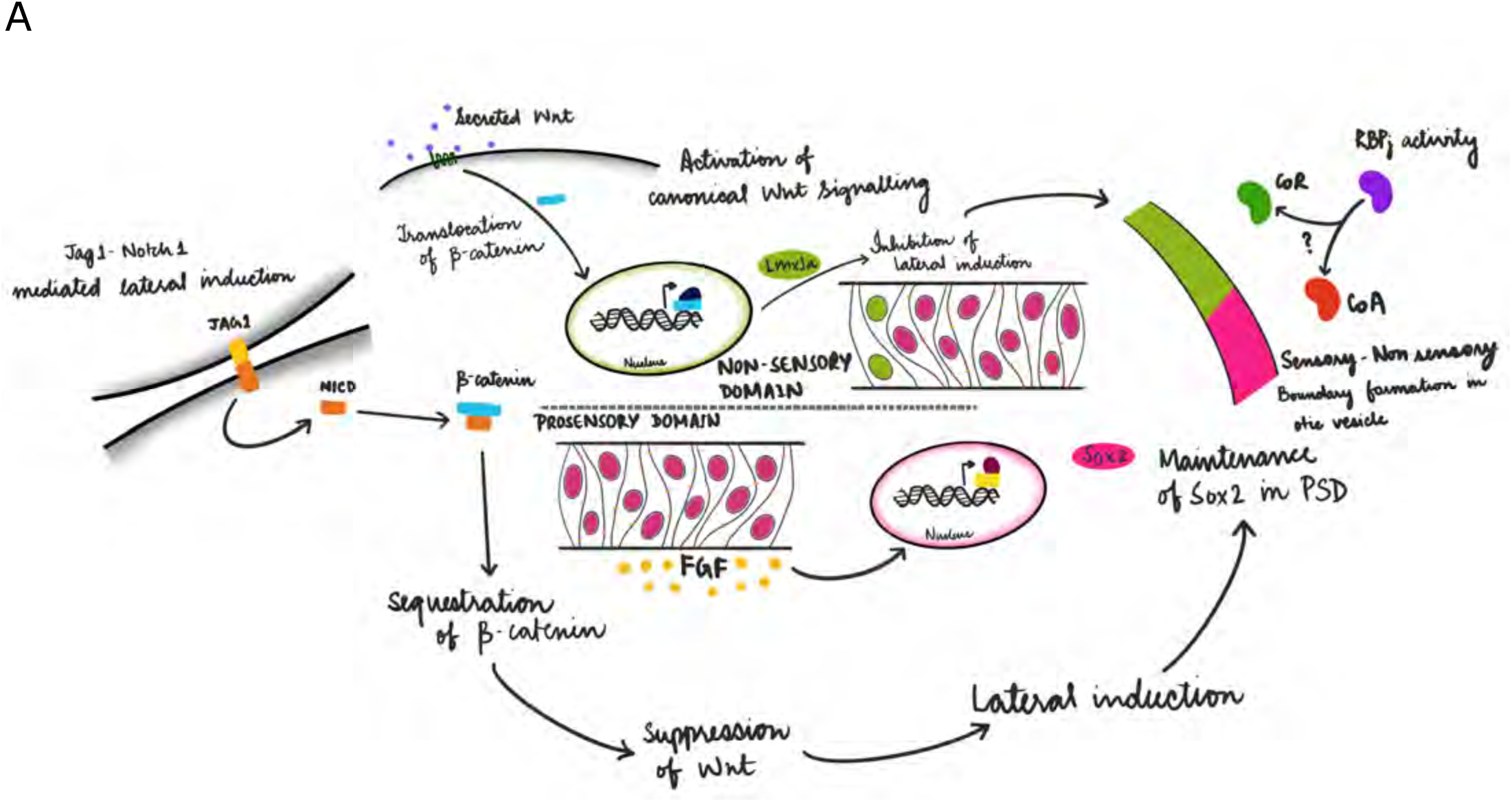
The WNTCH signalling pathway. (A) In the E9.5 embryo, Sox2 (magenta) expression becomes disrupted, and such cells are present throughout the otic vesicle, interspersed with Lmx1a-positive cells (green) in the double mutant embryo, β-catenin gain-of-function, and Jag1 cKO, instead of properly segregated sensory and non-sensory domains.

By approximately E14.0, the typical morphology of the inner ear, particularly the three semicircular canals, was evident in wild-type embryos. In contrast, the double mutants were difficult to interpret due to a severely disrupted morphology. This disruption is likely a consequence of excessive Wnt signalling activity throughout the otic vesicle (Fig. 7C).

## DISCUSSION

The formation of the prosensory domain (PSD) in the developing inner ear requires the coordinated action of multiple signalling pathways. Here, we show how Notch and Wnt signalling interact to establish and maintain the PSD, and how this interaction shapes the mutually exclusive expression of Sox2 and Lmx1a. Although Notch-mediated lateral induction has long been implicated in prosensory fate, the precise mechanism by which it maintains Sox2 expression has remained unresolved. We demonstrate that Jag1-mediated

Notch signalling is essential for maintaining, but not initiating, Sox2 in the PSD. Importantly, this regulation does not depend on canonical RBPjκ-dependent transcription, confirming a non-canonical mode of Notch activity during prosensory maintenance.

In contrast, Lmx1a, which marks non-sensory regions, remains unaffected in *Jag1* mutants but expands in *Rbpj*κ mutants. As RBPjκ can act as a transcriptional repressor by recruiting corepressors such as TLE and histone deacetylase complexes (Jennings and Ish-Horowicz, 2008), its loss may relieve repression, allowing Lmx1a to be ectopically expressed in regions where it is normally excluded. Thus, the emergence of Sox2/Lmx1a dual-positive domains in *Rbpj*κ mutants further suggests that boundary formation between sensory and non-sensory territories is not solely the result of mutually exclusive signalling leading to mutually exclusive transcriptional programs, but rather of these signals integrating to ensure lineage segregation.

We also identify Wnt signalling as a key upstream regulator of Lmx1a expression in both chick and mouse. Manipulating β-catenin levels shifts Lmx1a expression, confirming that Wnt functions as a dorsalising cue promoting non-sensory fate. Elevated Wnt activity represses Sox2, possibly through Lmx1a-mediated inhibition of Jag1-driven lateral induction, as reported in the chick (Mann et al., 2017). Conversely, complete loss of Wnt signalling via β-catenin deletion produces fragmented Sox2 domains, consistent with previous studies showing that moderate Wnt activity is necessary for proper prosensory specification (Zak and Daudet, 2021). These findings highlight that both excessive and absent Wnt signalling disrupt the precise spatial patterning of Sox2 expression.

A key finding of this study is that NICD and β-catenin interact directly in the cytoplasm. We propose that this interaction sequesters both proteins, preventing their nuclear accumulation. Cytoplasmic retention of β-catenin limits Wnt-dependent transcription and thereby stabilises Sox2 expression and prosensory identity. The implications of NICD sequestration are more complex. In the otocyst, Notch signalling operates in two overlapping but distinct modes: canonical Notch–Delta signalling drives neurogenesis in delaminating neuroblasts, while Jag1-mediated lateral induction maintains prosensory identity (Daudet and Zak, 2020; Kiernan, 2013). Retaining NICD in the cytoplasm within the prosensory domain could therefore suppress canonical Notch signalling, preventing neural differentiation while maintaining the prosensory state. β-catenin, in this context, acts not only as a Wnt effector but also as a buffering molecule that restricts NICD to the cytoplasm, ensuring appropriate spatio-temporal segregation of prosensory and neurogenic fates. Consistent with this, β- catenin loss-of-function in the chick leads to ectopic neurogenesis, suggesting that in its absence, NICD can translocate to the nucleus and activate neurogenic gene expression (Zak and Daudet, 2021). We propose that cytoplasmic NICD–β-catenin complexes function as a developmental checkpoint, preventing inappropriate neurogenic transcription within the PSD while neurogenesis proceeds in adjacent domains.

Together, these findings support a model in which Notch and Wnt signalling form an integrated regulatory module—a “WNTCH” axis (Muñoz-Descalzo et al., 2012). In this model, the PSD is maintained by a non-canonical Notch pathway in which NICD remains cytoplasmic, while dorsal non-sensory domains are patterned by β-catenin–dependent activation of Lmx1a. At their interface, NICD–β-catenin complexes act as a regulatory gate that balances prosensory stability with neurogenic potential. Disruption of either pathway shifts this balance, leading to sensory domain misspecification, loss of boundary integrity, or collapse of overall inner ear architecture. Thus, Notch and Wnt do not act as independent patterning cues, but as interdependent signalling partners whose dynamic interplay is essential for the coordinated establishment of sensory, neurogenic, and non-sensory identities in the early inner ear.

## ACKNOWLEDGMENT

This work was supported by the Department of Atomic Energy, Government of India, Project Identification No. RTI 4006, and grants from SERB, TIFR Infosys-Leading Edge Grant, the Royal National Institute for Deaf People. We acknowledge the support of the Animal Care and Resources Centre and the Central Imaging Facility at NCBS. We also acknowledge CEAH, Hessaraghatta, for providing us with fertilised eggs. We thank Prof. Nicholas Daudet for providing the Hes5- and TCF-reporters used for electroporation. We also thank Prof. Apurva Sarin for providing the NICD-NLS and NICD-NES plasmids. We acknowledge Prof. Shubha Tole and Prof. Makoto Taketo for β-catenin loss-of-function and gain-of-function mice. Finally, we thank all members of the Ear Lab at NCBS for discussions and valuable feedback.

---

**SUPPLEMENTARY FIGURE 1:** Expression of Notch signalling components in the otic vesicle. (A) Co-expression of Jag1 and Sox2 in the prosensory domain of the E11.5 inner ear. (B) Immunofluorescence to show the localisation of RBPjκ in the nuclei of the cells. (C) Expression of Sox2 is retained in the sensory epithelium despite the loss of *Rbpj*κ in the E11.5 embryo transverse section. Scale bar is 50μm

**SUPPLEMENTARY FIGURE 2:** Western blot membrane for the interaction of NICD and β-catenin HEK293T cells. (A) Full western blot membrane where the lanes have NICDNES-EGFP expressing HEK293T cells, negative control and the sample input for co-immunoprecipitation.

**SUPPLEMENTARY FIGURE 3:** Disrupting Jag1 and overactivating the Wnt signalling leads to complete abrogation of Dorso-Ventral polarity. (A) Expression of the Sox2 positive and interspersed Lmx1a positive cells throughout the otic vesicle in β-cateninin gain-of-function and Jag1 heterozygote floxed E9.5 embryo. Scale bar is 50 μm.

## Notes

### Competing Interest Statement

The authors have declared no competing interest.

## REFERENCES

Abello, G., Khatri, S., Giraldez, F. and Alsina, B. (2007). Early regionalization of the otic placode and its regulation by the Notch signaling pathway. Mech Dev 124, 631–645.

Adam, J., Myat, A., Le Roux, I., Eddison, M., Henrique, D., Ish-Horowicz, D. and Lewis, J. (1998). Cell fate choices and the expression of Notch, Delta and Serrate homologues in the chick inner ear: parallels with Drosophila sense-organ development. Development 125, 4645–4654.

An, W. F., Germain, A. R., Bishop, J. A., Nag, P. P., Metkar, S., Ketterman, J., Walk, M., Weiwer, M., Liu, X., Patnaik, D., et al. (2010). Discovery of Potent and Highly Selective Inhibitors of GSK3b. In Probe Reports from the NIH Molecular Libraries Program. Bethesda (MD).

Appler, J. M. and Goodrich, L. V. (2011). Connecting the ear to the brain: Molecular mechanisms of auditory circuit assembly. Prog Neurobiol 93, 488–508.

Basch, M. L., Ohyama, T., Segil, N. and Groves, A. K. (2011). Canonical Notch signaling is not necessary for prosensory induction in the mouse cochlea: insights from a conditional mutant of RBPjkappa. J Neurosci 31, 8046–8058.

Bray, S. J. (2006). Notch signalling: a simple pathway becomes complex. Nat Rev Mol Cell Biol 7, 678–689.

Brooker, R., Hozumi, K. and Lewis, J. (2006). Notch ligands with contrasting functions: Jagged1 and Delta1 in the mouse inner ear. Development 133, 1277–1286.

Chizhikov, V. V., Iskusnykh, I. Y., Fattakhov, N. and Fritzsch, B. (2021). Lmx1a and Lmx1b are Redundantly Required for the Development of Multiple Components of the Mammalian Auditory System. Neuroscience 452, 247–264.

Chung, S., Leung, A., Han, B. S., Chang, M. Y., Moon, J. I., Kim, C. H., Hong, S., Pruszak, J., Isacson, O. and Kim, K. S. (2009). Wnt1-lmx1a forms a novel autoregulatory loop and controls midbrain dopaminergic differentiation synergistically with the SHH-FoxA2 pathway. Cell stem cell 5, 646–658.

Cole, L. K., Le Roux, I., Nunes, F., Laufer, E., Lewis, J. and Wu, D. K. (2000). Sensory organ generation in the chicken inner ear: contributions of bone morphogenetic protein 4, serrate1, and lunatic fringe. J Comp Neurol 424, 509–520.

Dabdoub, A., Puligilla, C., Jones, J. M., Fritzsch, B., Cheah, K. S., Pevny, L. H. and Kelley, M. W. (2008). Sox2 signaling in prosensory domain specification and subsequent hair cell differentiation in the developing cochlea. Proc Natl Acad Sci U S A 105, 18396–18401.

Daudet, N., Ariza-McNaughton, L. and Lewis, J. (2007). Notch signalling is needed to maintain, but not to initiate, the formation of prosensory patches in the chick inner ear. Development 134, 2369–2378.

Daudet, N. and Zak, M. (2020). Notch Signalling: The Multitask Manager of Inner Ear Development and Regeneration. Advances in experimental medicine and biology 1218, 129–157.

Fekete, D. M. and Wu, D. K. (2002). Revisiting cell fate specification in the inner ear. Curr Opin Neurobiol 12, 35–42.

Groves, A. K. and Fekete, D. M. (2012). Shaping sound in space: the regulation of inner ear patterning. Development 139, 245–257.

Gu, R., Brown, R. M., 2nd, Hsu, C. W., Cai, T., Crowder, A. L., Piazza, V. G., Vadakkan, T. J., Dickinson, M. E. and Groves, A. K. (2016). Lineage tracing of Sox2-expressing progenitor cells in the mouse inner ear reveals a broad contribution to non-sensory tissues and insights into the origin of the organ of Corti. Dev Biol 414, 72–84.

Harada, N., Tamai, Y., Ishikawa, T., Sauer, B., Takaku, K., Oshima, M. and Taketo, M. M. (1999). Intestinal polyposis in mice with a dominant stable mutation of the beta-catenin gene. EMBO J 18, 5931–5942.

Hayashi, T., Cunningham, D. and Bermingham-McDonogh, O. (2007). Loss of Fgfr3 leads to excess hair cell development in the mouse organ of Corti. Dev Dyn 236, 525–533.

Huang, S. M., Mishina, Y. M., Liu, S., Cheung, A., Stegmeier, F., Michaud, G. A., Charlat, O., Wiellette, E., Zhang, Y., Wiessner, S., et al. (2009). Tankyrase inhibition stabilizes axin and antagonizes Wnt signalling. Nature 461, 614–620.

Jacques, B. E., Puligilla, C., Weichert, R. M., Ferrer-Vaquer, A., Hadjantonakis, A. K., Kelley, M. W. and Dabdoub, A. (2012). A dual function for canonical Wnt/beta-catenin signaling in the developing mammalian cochlea. Development 139, 4395–4404.

Jennings, B. H. and Ish-Horowicz, D. (2008). The Groucho/TLE/Grg family of transcriptional co-repressors. Genome Biol 9, 205.

Kiernan, A. E. (2013). Notch signaling during cell fate determination in the inner ear. Semin Cell Dev Biol 24, 470–479.

Kiernan, A. E., Cordes, R., Kopan, R., Gossler, A. and Gridley, T. (2005). The Notch ligands DLL1 and JAG2 act synergistically to regulate hair cell development in the mammalian inner ear. Development 132, 4353–4362.

Kiernan, A. E., Xu, J. and Gridley, T. (2006). The Notch Ligand JAG1 Is Required for Sensory Progenitor Development in the Mammalian Inner Ear. PLoS Genet 2, e4.

Koo, S. K., Hill, J. K., Hwang, C. H., Lin, Z. S., Millen, K. J. and Wu, D. K. (2009). Lmx1a maintains proper neurogenic, sensory, and non-sensory domains in the mammalian inner ear. Dev Biol 333, 14–25.

Ladher, R. K., O’Neill, P. and Begbie, J. (2010). From shared lineage to distinct functions: the development of the inner ear and epibranchial placodes. Development 137, 1777–1785.

Mann, Z. F., Galvez, H., Pedreno, D., Chen, Z., Chrysostomou, E., Zak, M., Kang, M., Canden, E. and Daudet, N. (2017). Shaping of inner ear sensory organs through antagonistic interactions between Notch signalling and Lmx1a. eLife 6.

Muñoz Descalzo, S., de Navascues, J., & Arias, A. M. (2012). Wnt Notch signalling: An integrated mechanism regulating transitions between cell states. Bioessays, 34(2), 110–118.

Nelson, J. C., Hosamani, I. V. and Groves, A. K. (2025). Control of sensory cell differentiation in the inner ear by extracellular signals and transcriptional regulators. Curr Top Dev Biol 165, 1–44.

Neves, J., Parada, C., Chamizo, M. and Giraldez, F. (2011). Jagged 1 regulates the restriction of Sox2 expression in the developing chicken inner ear: a mechanism for sensory organ specification. Development 138, 735–744.

Noda, T., Oki, S., Kitajima, K., Harada, T., Komune, S. and Meno, C. (2012). Restriction of Wnt signaling in the dorsal otocyst determines semicircular canal formation in the mouse embryo. Dev Biol 362, 83–93.

Ohyama, T., Basch, M. L., Mishina, Y., Lyons, K. M., Segil, N. and Groves, A. K. (2010). BMP signaling is necessary for patterning the sensory and nonsensory regions of the developing mammalian cochlea. J Neurosci 30, 15044–15051.

Ono, K., Kita, T., Sato, S., O’Neill, P., Mak, S. S., Paschaki, M., Ito, M., Gotoh, N., Kawakami, K., Sasai, Y., et al. (2014). FGFR1-Frs2/3 signalling maintains sensory progenitors during inner ear hair cell formation. PLoS Genet 10, e1004118.

Oswald, F. and Kovall, R. A. (2018). CSL-Associated Corepressor and Coactivator Complexes. Advances in experimental medicine and biology 1066, 279–295.

Raft, S., Koundakjian, E. J., Quinones, H., Jayasena, C. S., Goodrich, L. V., Johnson, J. E., Segil, N. and Groves, A. K. (2007). Cross-regulation of Ngn1 and Math1 coordinates the production of neurons and sensory hair cells during inner ear development. Development 134, 4405–4415.

Riccomagno, M. M., Martinu, L., Mulheisen, M., Wu, D. K. and Epstein, D. J. (2002). Specification of the mammalian cochlea is dependent on Sonic hedgehog. Genes Dev 16, 2365–2378.

Riccomagno, M. M., Takada, S. and Epstein, D. J. (2005). Wnt-dependent regulation of inner ear morphogenesis is balanced by the opposing and supporting roles of Shh. Genes Dev 19, 1612–1623.

Sai, X. and Ladher, R. K. (2015). Early steps in inner ear development: induction and morphogenesis of the otic placode. Front Pharmacol 6, 19.

Sato, S., Furuta, Y. and Kawakami, K. (2018). Regulation of continuous but complex expression pattern of Six1 during early sensory development. Dev Dyn 247, 250–261.

Tamilkumar, V. N., Purushothama, H. and Ladher, R. K. (2025). Epithelial fusion is mediated by a partial epithelial-mesenchymal transition. Biology open 14.

Wright, T. J. and Mansour, S. L. (2003). FGF signaling in ear development and innervation. Curr Top Dev Biol 57, 225–259.

Wu, D. K. and Kelley, M. W. (2012). Molecular mechanisms of inner ear development. Cold Spring Harbor perspectives in biology 4, a008409.

Yamamoto, N., Chang, W. and Kelley, M. W. (2011). Rbpj regulates development of prosensory cells in the mammalian inner ear. Dev Biol 353, 367–379.

Zak, M. and Daudet, N. (2021). A gradient of Wnt activity positions the neurosensory domains of the inner ear. eLife 10.

