## Supplementary for "Notch and Wnt signalling interact for proper prosensory and non-sensory domain formation"

### SUPPLEMENTARY FIGURE 1

Expression of Notch signalling components in the otic epithelium

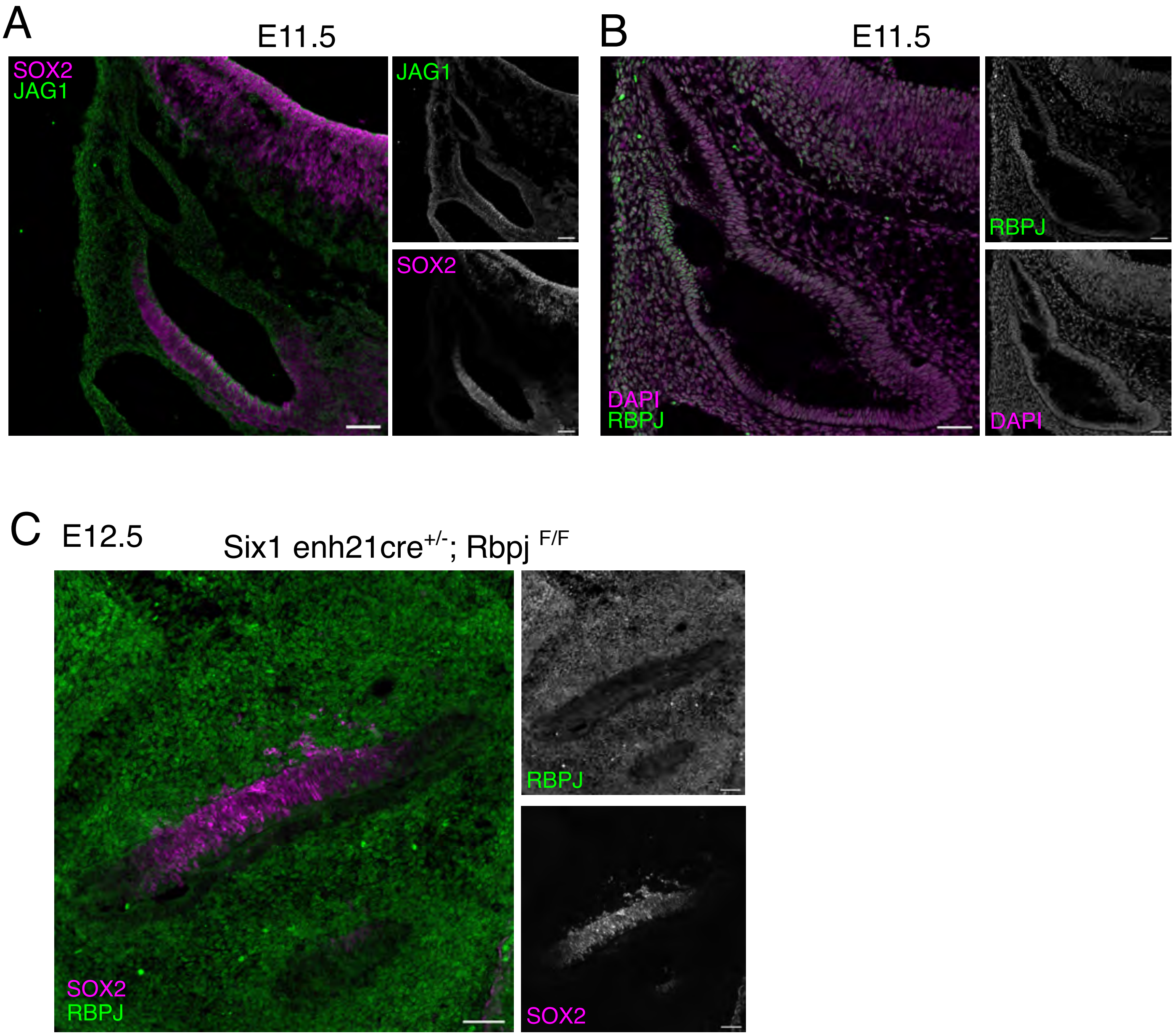

### SUPPLEMENTARY FIGURE 2

Western blot membrane for the interaction of NICD and β-catenin in HEK293

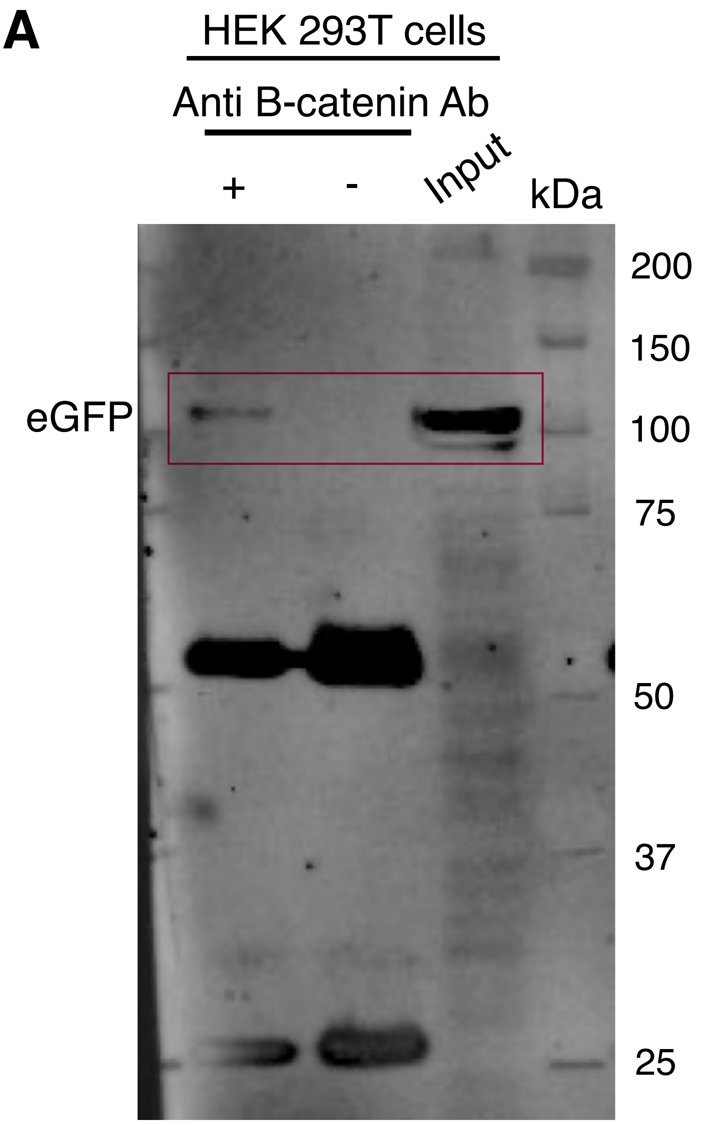

### SUPPLEMENTARY FIGURE 3

Disrupting Jag1 and over activating the Wnt signalling leads to complete abrogation of Dorso-Ventral polarity

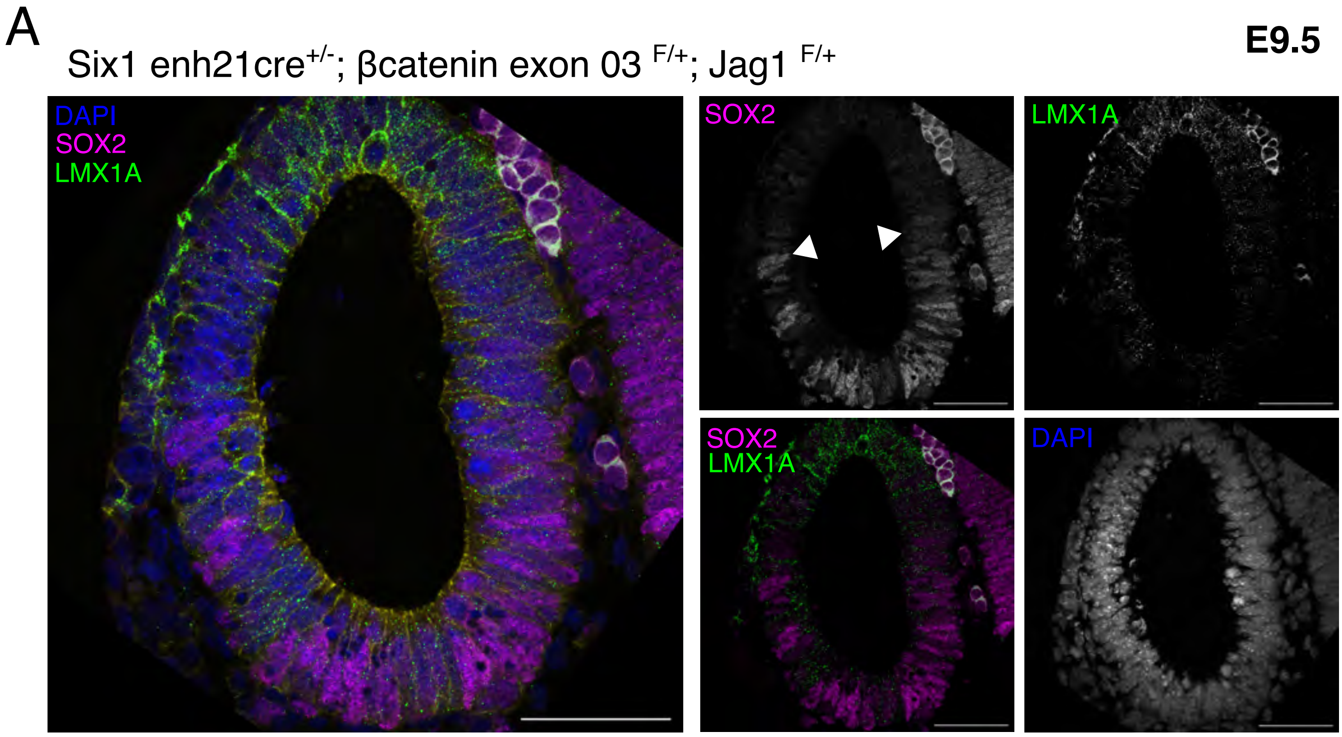
